# scMaize: A Single-Cell Foundation Model and Integrated Atlas for Maize

**DOI:** 10.64898/2026.08.01.742180

**Authors:** Qian Cheng, Ying Zhang, Tianhao Wu, Anwen Zhao, Meiqi Shang, Xiangfeng Wang, Jun Yan

## Abstract

Single-cell transcriptomics has resolved cell-type-specific gene expression in plants, yet maize still lacks an integrated reference and species-specific foundation models. We present scMaize, combining scMaizeAtlas, an integrated atlas of 385,675 cells from 20 projects and 66 samples across seven tissues with hierarchical annotation, with two Transformer-based foundation models pretrained on this atlas. scMaizeExp serves as an expression-only baseline, while scMaizeGO incorporates Gene Ontology (GO) functional embeddings as an inductive bias. Although global expression-prediction accuracy was comparable, the GO prior improved rank-order prediction, strengthened attention toward functionally coherent gene modules, and enhanced embedding topology, with scMaizeGO achieving 86.0% cell-type and 97.1% tissue classification accuracy. Zero-shot evaluation demonstrated the cross-species generalizability of scMaizeGO representations, and few-shot fine-tuning enabled accurate cross-species classification with minimal labeled data. Root perturbation-condition analysis showed that the model encoded treatment-specific cellular states beyond cell-type identity, with the GO prior amplifying perturbation signals approximately threefold. Expression projection identified condition-responsive genes enriched for known stress pathways, and attention analysis revealed predominantly condition-specific changes in gene-gene attention that were weakly associated with expression-projection changes. An online platform (https://www.scmaize.com) provides atlas exploration, model access, and zero-code analysis tools. scMaize establishes a framework demonstrating that species-specific pretraining with functional priors enables transferable, perturbation-aware representations for crop single-cell genomics.

**HIGHLIGHTS:**

- scMaizeAtlas integrates 385,675 cells from 20 maize single-cell projects.
- scMaizeGO incorporates Gene Ontology priors into maize-specific pretraining.
- GO priors improve rank-order prediction, attention coherence and embeddings.
- Few-shot tuning enables cross-species cell-type classification with limited labels.
- Expression projection reveals stress-responsive genes in root cell states.

## INTRODUCTION

Single-cell genomics and artificial intelligence are reshaping the resolution at which plant development, environmental adaptation and agronomic variation can be interrogated (Yan and Wang, 2023). Single-cell RNA sequencing (scRNA-seq) resolves gene expression programs within defined cell populations, revealing signals that are diluted or obscured in bulk tissue profiles (Grones et al., 2024; Szalata et al., 2024). In plants, the field has progressed from single-organ surveys to organism-scale and cross-tissue references, including recent atlases for *Arabidopsis*, rice, soybean and vascular plants more broadly (Guo et al., 2025; Wang et al., 2025b; Xue et al., 2025; Zhang et al., 2025). These resources have established cell-type-resolved frameworks for marker discovery, regulatory inference and candidate gene prioritization.

Maize (*Zea mays* L.) is both a major crop species and a long-standing model for genetics, development and breeding. Multiple single-cell datasets have been generated from roots, leaves, reproductive tissues and developing seeds (Marand et al., 2021; Marand et al., 2025; Ortiz-Ramírez et al., 2021; Wang et al., 2025a; Yuan et al., 2024). However, these datasets remain partitioned by study, platform, genotype, developmental stage and processing pipeline. Consequently, the maize community lacks a unified reference in which cells from diverse tissues and studies can be compared within a consistent coordinate system. This fragmentation also constrains model development, because algorithms trained on isolated datasets tend to capture study-specific structure rather than reusable maize expression programs.

In single-cell biology, scBERT (Yang et al., 2022), Geneformer (Theodoris et al., 2023), scGPT (Cui et al., 2024), and scFoundation (Hao et al., 2024a) have shown that Transformer architectures trained on millions of transcriptomes encode biologically meaningful relationships among cells and genes, as highlighted in a recent review on foundation models (Xu et al., 2025). The emergence of cross-species foundation models like GeneCompass (Yang et al., 2024) and TranscriptFormer (Pearce et al., 2026) further underscores the value of large-scale corpora and structured biological priors for decoding general gene regulatory principles. Meanwhile, plant-dedicated models such as scPlantLLM (Cao et al., 2025) and scPlantFormer (Zhang et al., 2024), though smaller in scale and lacking explicit prior integration, suggest that domain-specific training data can also benefit plant single-cell analysis. Nevertheless, it remains unclear whether species-focused pretraining on maize data can capture organism-specific expression architecture more effectively than broadly transferred representations.

Functional annotation provides a complementary source of biological structure. The Gene Ontology (GO) organizes curated knowledge of molecular functions, biological processes and cellular components (Gene Ontology Consortium, 2021). Rather than forcing a model to infer all functional relationships from sparse expression matrices, GO-derived embeddings can impose an interpretable inductive bias on representation learning. Human single-cell models have begun to exploit GO information (Bai et al., 2026; Zeng et al., 2025), but its utility in plant single-cell foundation modeling has not been systematically tested.

Here we introduce scMaize, an integrated maize single-cell atlas and species-specific foundation-model framework. We harmonized over 0.6 million initially collected cells from 20 studies into 385,675 high-quality cells and developed a hierarchical annotation system for maize cell identities. We then trained two Transformer models, scMaizeExp and scMaizeGO, and assessed them through expression prediction, GO-guided attention analysis, hub-gene enrichment and embedding benchmarks. To assess the generality of these learned representations, we conducted external validation on three independent datasets spanning maize, rice and *Arabidopsis*. Few-shot fine-tuning on these datasets achieved accurate cross-dataset and cross-species cell-type classification with minimal labeled data. We further investigated whether the model captures cellular states beyond cell-type identity by examining its embedding dynamics under perturbation conditions, and evaluated expression projection and gene–gene attention changes as complementary readouts for perturbation-aware gene discovery. Finally, we deployed an online platform (https://www.scmaize.com) that provides interactive atlas exploration, model access and zero-code analysis tools to support community adoption.

## RESULTS

### An integrated single-cell atlas of maize cellular diversity

To construct a reference for maize cellular diversity, we collected scRNA-seq data from 20 public projects comprising 66 samples across seven tissue systems: seedling, leaf, root, shoot apex, ear, tassel and endosperm (Figure 1A). The collection spans B73, additional inbred lines and hybrids, and includes untreated samples, stress conditions and multiple developmental stages. A total of 681,032 initially collected cells were subjected to uniform quality control and filtering, yielding 385,675 high-quality cells for integration. Sample-aware batch correction was applied, and UMAP visualization before and after correction demonstrated that technical variation was substantially reduced while tissue-level structure was preserved (Figure 1B). Leiden clustering resolved 30 transcriptionally distinct clusters (clusters 0–29; Figure 1C). The post-annotation organization showed that cell types shared across tissues tended to co-cluster, whereas tissue-restricted populations remained distinct, indicating that the integrated atlas was structured primarily by cell identity rather than project origin.

**Figure 1.**
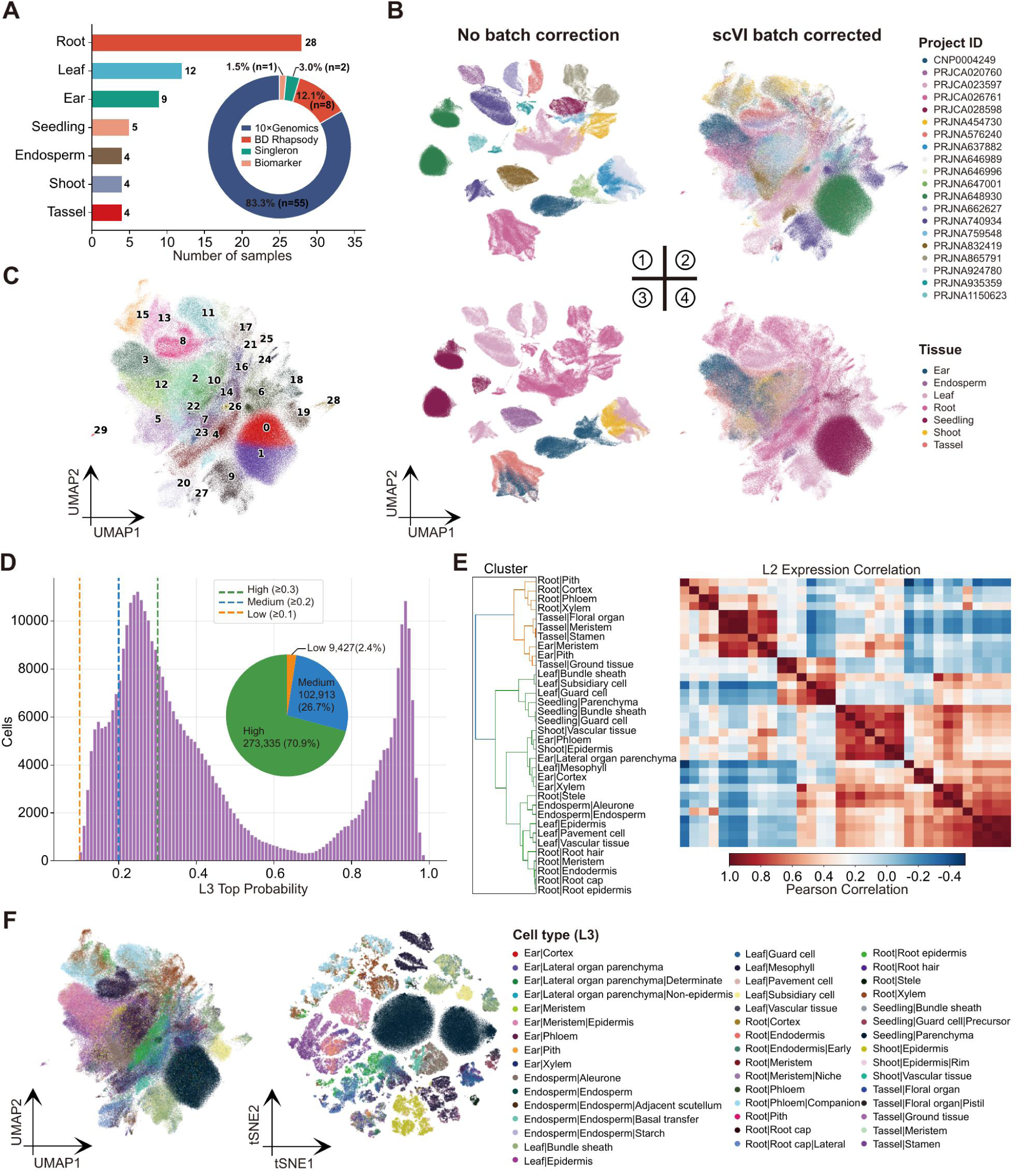
Construction and annotation of the scMaizeAtlas. (A) Summary of the 20 integrated projects, including sample number, tissue coverage and sequencing-platform composition. (B) UMAP visualization before and after scVI batch correction, colored by project of origin and tissue type. (C) Leiden clustering of the integrated atlas, showing 30 clusters (0–29). (D) Annotation confidence distribution, including histogram and pie chart of High, Medium and Low confidence assignments. (E) Pearson correlation heatmap of mean marker expression across L2 cell types. (F) UMAP and t-SNE visualizations colored by L3 cell-type annotation, showing discrete lineages and partially continuous cell states.

Cluster annotation required a marker system resilient to noise and inconsistent nomenclature in existing plant resources. We curated 70,023 marker records from scPlantDB (He et al., 2024), PlantscRNAdb (Chen et al., 2021) and published maize single-cell studies. Because 72% of genes were assigned as markers for multiple cell types without statistical support, we applied a refinement procedure involving statistical validation against the integrated atlas, specificity filtering, GO embedding-based deduplication and hierarchical mapping. This process produced 3,074 non-redundant markers assigned to 34 L2 and 46 L3 categories. Annotation confidence remained high across the hierarchy: 77.4% of cells were assigned at L3 resolution, and 70.9% of cells received high-confidence labels (Figure 1D and Supplementary Figure S1). Pearson correlation of marker expression among L2 types recovered expected relationships, including the grouping of meristematic cell types from tassel and ear, epidermal subtypes across tissues and a distinct vascular block (Figure 1E). By contrast, different cell types within the same tissue showed divergent expression profiles. UMAP and t-SNE views of cell types revealed both discrete lineages and partially overlapping states, consistent with the coexistence of stable identities and developmental continua across the atlas (Figure 1F).

### A species-specific Transformer model for maize single-cell data

We designed a maize-focused Transformer encoder to learn cell and gene representations from the integrated atlas (Figure 2A). Each cell was represented by a compact sequence of 2,048 genes sampled from the 15,000-gene vocabulary and ordered by highly variable gene (HVG) rank. This strategy exposes the model to diverse gene combinations while retaining a sequence length compatible with efficient GPU training. The architecture includes a learnable CLS token for cell-level representation and four input branches encoding gene identity, continuous expression values, batch labels and, in scMaizeGO, GO functional embeddings. The core encoder contains 6 Transformer layers with hidden dimension 384, 4 attention heads and feed-forward dimension 1,536. scMaizeExp and scMaizeGO contain 16.7M and 16.8M parameters, respectively.

**Figure 2.**
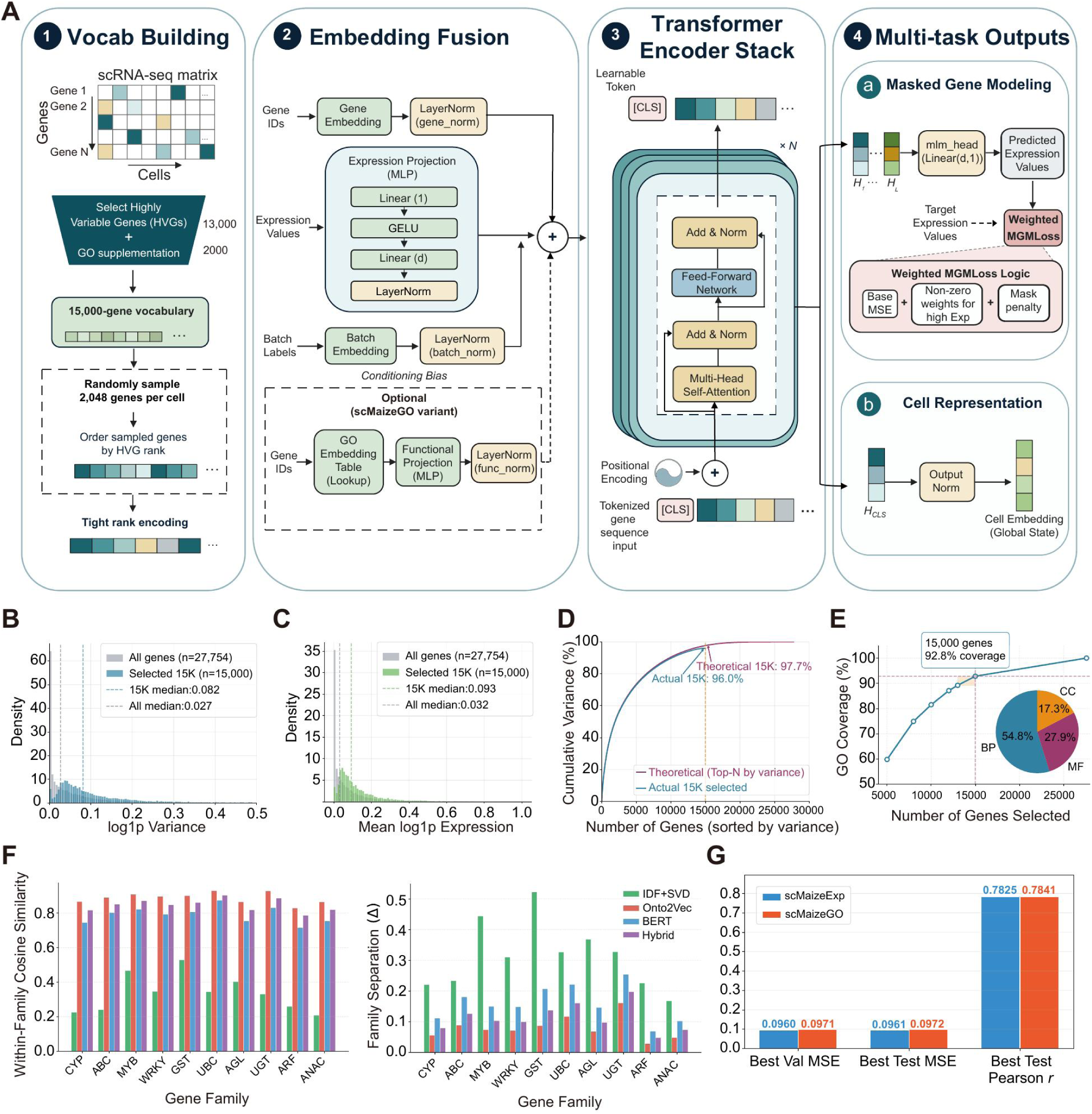
Design and training of scMaize foundation models. (A) scMaize Transformer architecture, including vocabulary construction, gene identity embedding, expression projection, batch conditioning, optional GO functional embedding, Transformer encoder layers, masked gene modeling output and CLS-based cell representation. (B,C) Distribution of all detected genes and the selected 15,000-gene vocabulary by expression variance (B) and mean expression (C). (D) Cumulative variance captured by the selected 15,000 genes compared with the theoretical top-variance selection. (E) GO coverage as a function of selected gene number and final BP/MF/CC composition. (F) GO embedding quality assessed by within-family and between-family cosine similarity for the top-10 Pfam families. (G) Validation and test performance of both models.

The vocabulary was built from 13,000 HVGs supplemented with 2,000 GO-coverage genes selected to maximize functional-category representation (Figure 2B–E). Among four evaluated GO embedding strategies, IDF-weighted singular value decomposition (IDF+SVD) achieved the largest family-separation score (0.278; Figure 2F and Supplementary Figure S2), and was therefore used for scMaizeGO. Both models were pretrained with 15% masking and a weighted MSE loss that upweighted errors on non-zero expression values. Following pretraining, the two models achieved nearly identical test Pearson correlations (scMaizeExp, 0.7825; scMaizeGO, 0.7841; Figure 2G), indicating that GO information did not materially alter global expression-prediction accuracy. The pretrained model supports masked gene prediction for expression imputation and gene-context analysis, while the CLS token provides a 384-dimensional cell embedding suitable for clustering, classification, visualization and other downstream learning tasks.

### Expression prediction is constrained by gene abundance and variability

We examined whether prediction accuracy was associated with intrinsic gene properties by stratifying genes according to mean expression and expression variance. Global performance was similar for scMaizeExp and scMaizeGO, with near-identical per-gene Pearson distributions (Figure 3A,E). Stratification revealed pronounced heterogeneity. Highly expressed genes were predicted more accurately than low-expression genes, and expression variance had an even stronger effect (Kruskal-Wallis *P* < 10^−300^ for both factors). The expression-by-variance heatmaps showed a monotonic gradient: genes with high expression and high variance were predicted most accurately, whereas genes with low expression and low variance remained the most difficult (Figure 3B–D,F–H). Transcription factors, which are often weakly expressed, were concentrated in this difficult region for both models, highlighting a limitation of expression-only prediction for regulatory genes. Despite comparable global accuracy, scMaizeGO showed stronger rank-order preservation (Figure 3I). The GO prior therefore improved the model’s ability to recover relative expression differences rather than absolute values, a property relevant for differential expression analysis.

**Figure 3.**
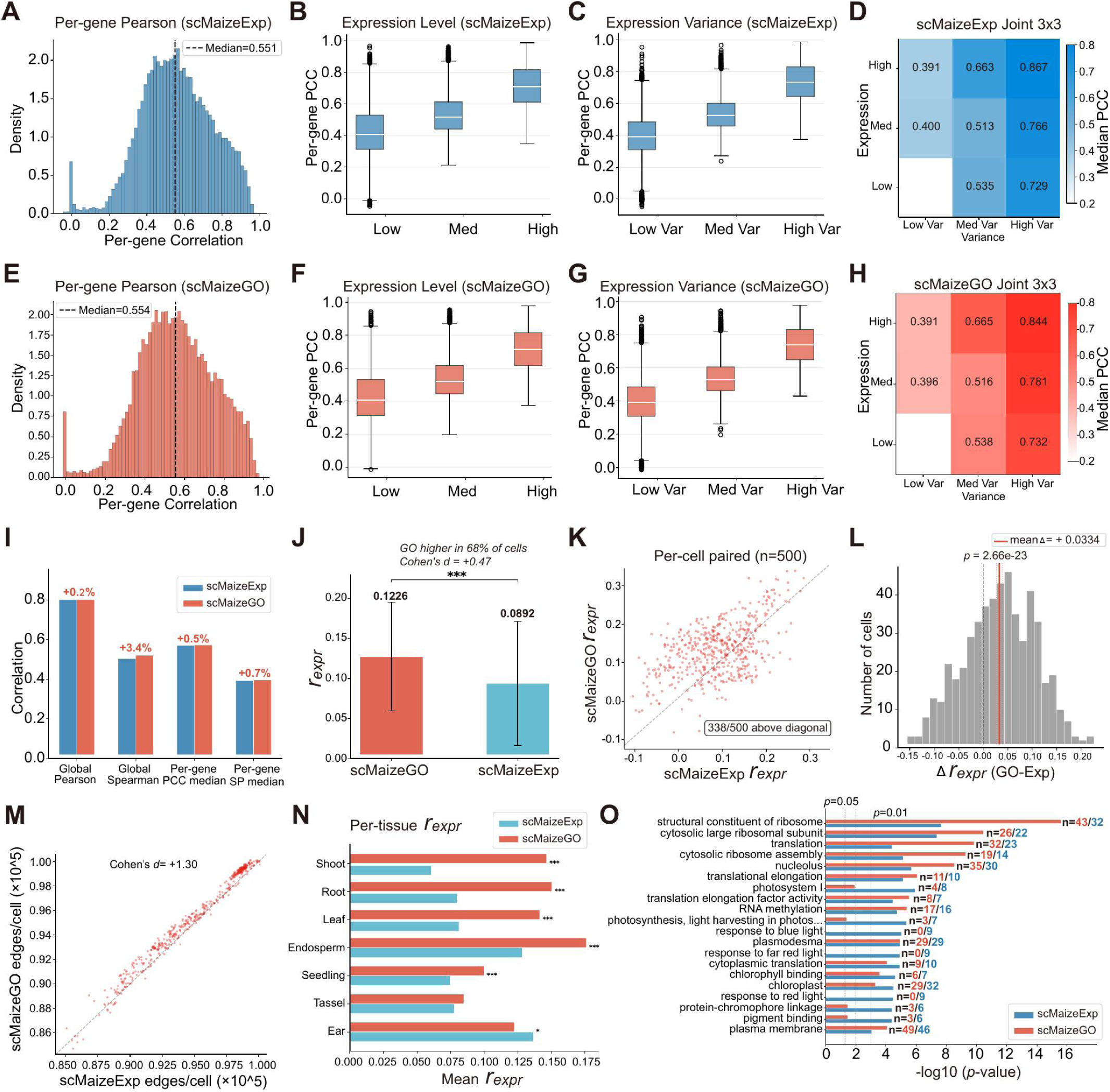
Expression prediction and attention mechanism analysis. (A–D) scMaizeExp per-gene Pearson distribution, expression-level stratification, variance stratification and joint 3 × 3 heatmap. (E–H) Corresponding analyses for scMaizeGO. (I) Global and per-gene Pearson and Spearman correlations, with relative changes between models. (J) Bar plot of rexpr for both models. (K) Paired scatter plot of per-cell rexpr for scMaizeGO versus scMaizeExp based on 500 randomly selected cells. (L) Histogram of the per-cell difference Δrexpr (scMaizeGO − scMaizeExp). (M) Paired comparison of attention edge counts. (N) Tissue-specific rexpr effects. (O) Functional enrichment of hub genes identified from cumulative outgoing attention.

### GO prior strengthens attention toward co-expressed and functionally dedicated genes

The attention mechanism provided insight into how each model distributed its focus across gene pairs. The critical differentiator was the alignment between attention and gene co-expression structure. scMaizeGO consistently exhibited higher *r*_expr_ — the Spearman correlation between per-cell pairwise attention and expression similarity — than scMaizeExp (0.1226 versus 0.0892; Cohen’s *d* = 0.47; Figure 3J). Paired analysis of 500 cells showed that 338 cells (67.6%) had higher *r*_expr_ in scMaizeGO (Figure 3K), with a mean Δ*r*_expr_ (GO − Exp) of 0.0334 (*P* = 2.66 × 10^−23^; Figure 3L). This integration of GO prior regularized the attention landscape, concentrating attention onto gene pairs whose expression was coupled across cells. The difference was also evident at the gene-pair level: scMaizeGO produced more attention edges than scMaizeExp (Cohen’s *d* = 1.30; Figure 3M), suggesting that the GO prior contributed additional gene-gene associations beyond those captured by expression alone. The *r*_expr_ advantage was most pronounced in tissues with complex cell-type compositions, such as root and shoot apex, where intercellular expression heterogeneity is high (Figure 3N). This tissue-specific pattern suggests that GO information may be particularly informative in heterogeneous cellular environments, where expression-based signals alone are less decisive for resolving biologically meaningful gene-gene relationships.

Hub genes — the top-100 genes ranked by outgoing attention — differed between the two models. scMaizeGO hub genes were enriched for basic cellular machinery: ribosome, translation and nucleolus. scMaizeExp hubs showed comparable enrichment for these categories but with lower fold enrichment scores (Figure 3O), and additional signals emerged for photosynthesis pathways. Both models underrepresented transcription factors among attention hubs. This observation aligns with findings from single-cell foundation models, where attention patterns are known to be strongly influenced by gene expression levels (Kendiukhov, 2026), often biasing toward highly expressed housekeeping genes (Theodoris et al., 2023).

### scMaize embeddings improve cell-type discrimination and manifold coherence

To evaluate the practical value of scMaize embeddings for cell representation, CLS embeddings from 27,365 high-confidence test cells were compared with PCA30_HVG (PCA on highly variable genes, without batch correction), Harmony_HVG (Harmony-corrected), and scPlantLLM embeddings, including both general pretrained and maize fine-tuned versions. UMAPs colored by tissue and by L2 cell type showed that both scMaize models produced coherent manifolds, with related cell types occupying adjacent regions and tissue identity appearing as continuous gradients rather than isolated blocks (Figure 4A,B). This behavior was most evident for scMaizeGO. By contrast, PCA30_HVG embeddings were dominated by batch effects, which obscured tissue-specific structures and confounded cell-type distinctions. Harmony_HVG, on the other hand, compressed several distinct cell types into overlapping regions, consistent with over-harmonization under strong batch correction, whereas scPlantLLM embeddings appeared fragmented and less organized in the maize test setting.

**Figure 4.**
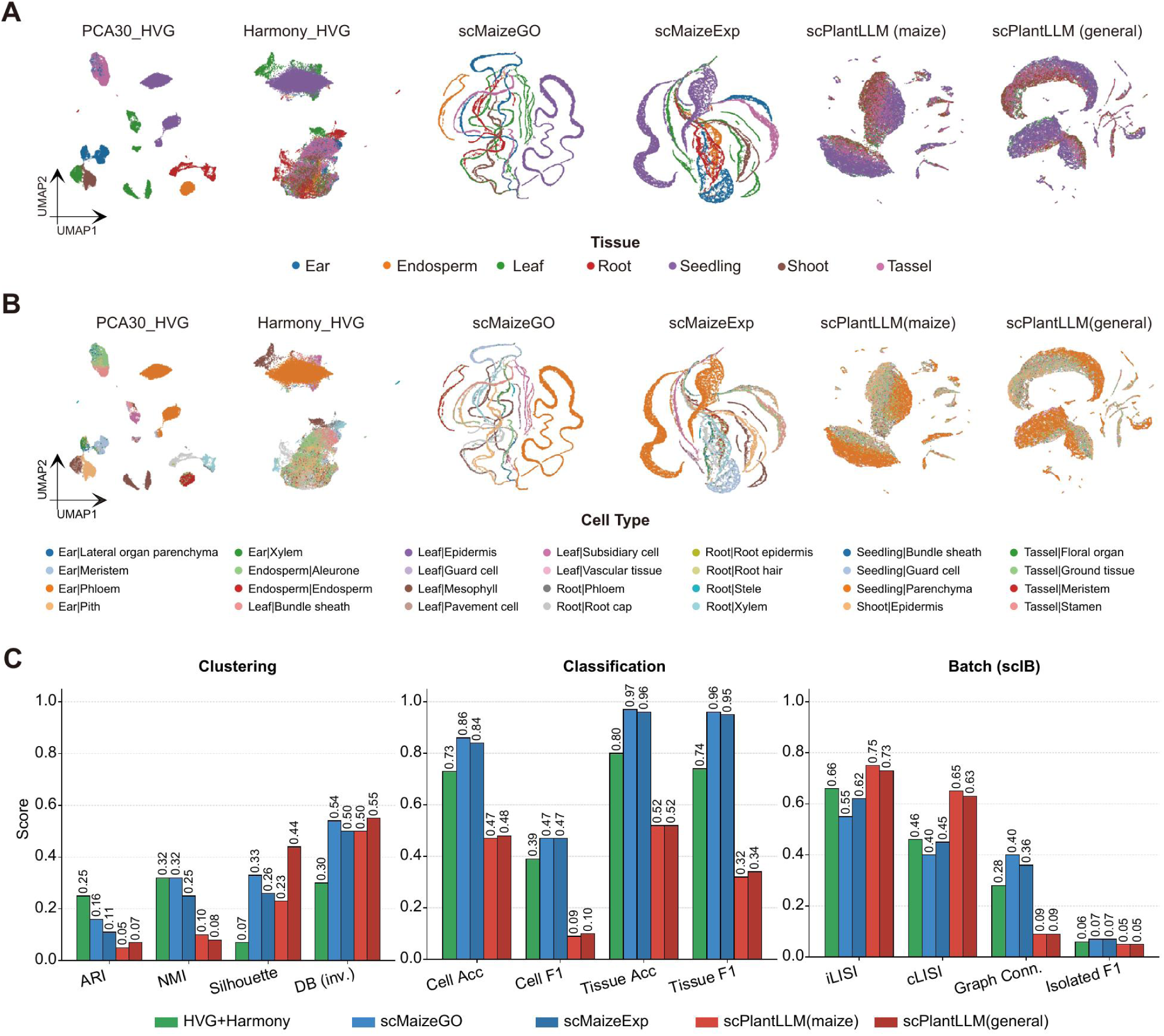
Cell embedding evaluation and downstream applications. (A) UMAP visualization of embeddings colored by tissue for PCA30_HVG, Harmony_HVG, scMaizeGO, scMaizeExp, scPlantLLM maize fine-tuned and scPlantLLM general models. (B) The same embeddings colored by L2 cell type. (C) Quantitative benchmark across clustering, classification and batch-integration metrics, including ARI, NMI, silhouette score, inverse Davies-Bouldin index, cell-type and tissue classification metrics, iLISI, cLISI, graph connectivity and isolated-label F1.

Figure 4C quantitatively compared Harmony_HVG and the four model embeddings. Across twelve metrics covering clustering, classification and batch integration, scMaizeGO achieved 86.0% cell-type classification accuracy and 97.1% tissue classification accuracy, compared with 72.9% and 79.7% for Harmony_HVG and 46.8% and 51.7% for the maize fine-tuned scPlantLLM. Although the scMaize models were not designed as explicit batch-correction algorithms, scMaizeGO showed higher graph connectivity than Harmony_HVG (0.397 versus 0.275; a measure of within-type neighborhood connectivity), indicating better preservation of biological neighborhood structure. Its iLISI was lower than that of Harmony_HVG (0.551 versus 0.664), reflecting a trade-off between preserving biological signal and achieving maximum batch mixing — an expected outcome given that scMaize uses batch labels as conditioning variables rather than as an optimization target for full batch removal. For clustering interpretation, we emphasize ARI and NMI. The lower ARI of scMaizeGO relative to Harmony_HVG likely reflects the preservation of continuous cell-state gradients within the same cell-type label, which discrete clustering metrics such as ARI are not designed to capture. This interpretation is consistent with the observation that ARI, as a label-based metric, penalizes deviations from discrete cluster assignments even when such deviations represent genuine biological continuity.

### Cross-species generalization of learned cell representations

To assess cross-dataset and cross-species transferability, we performed zero-shot cell-type classification on three external datasets spanning maize root (15,901 cells), rice root tip (26,138 cells) and *Arabidopsis* leaf (5,594 cells), with rice and *Arabidopsis* genes mapped to maize orthologs via one-to-one orthology relationships (∼79% and ∼70% gene coverage). Zero-shot evaluation showed that scMaizeGO consistently outperformed scMaizeExp across all three datasets (Figure 5A). On maize root, scMaizeGO achieved a k-NN macro F1 score of 0.292 versus 0.249 for scMaizeExp; on rice root, 0.398 versus 0.236; and on *Arabidopsis* leaf, 0.553 versus 0.471. Across all species, scMaizeGO embeddings exhibited lower effective dimensionality with comparable or higher neighbor consistency, indicating more compact and biologically coherent representations. Examination of the raw embedding variance revealed that scMaizeGO compressed cellular information into ∼30–40 effective dimensions, with principal component analysis (PCA) further reducing this to fewer than 10 dimensions (Figure 5B). This compactness reflects the representation-learning capacity of masked gene modeling, which distills high-dimensional expression data into low-dimensional features. However, such dimensional concentration may also limit the resolution of fine-grained cell-type distinctions.

**Figure 5.**
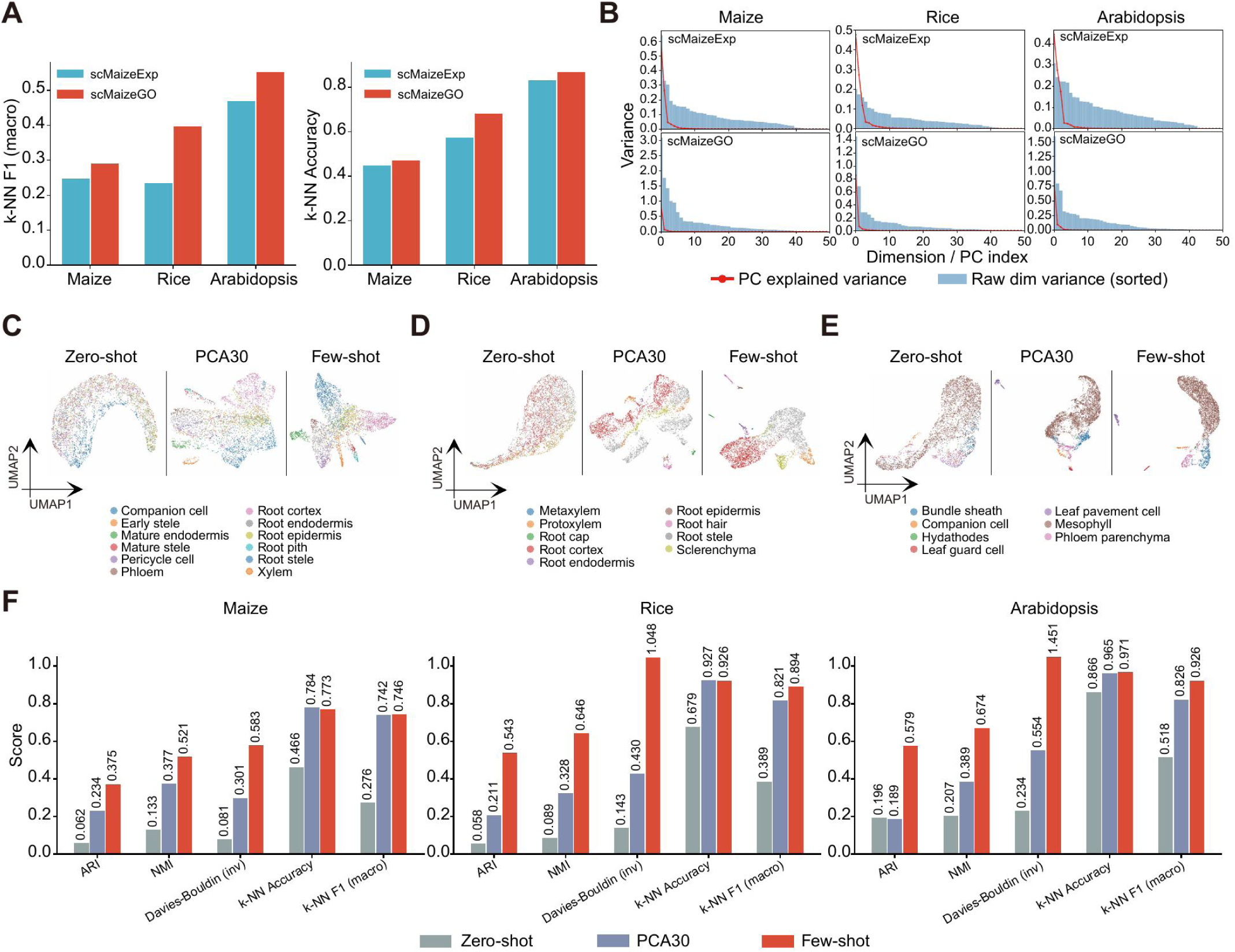
External validation and few-shot fine-tuning of scMaize embeddings. (A) Zero-shot k-NN macro F1 and accuracy of scMaizeExp and scMaizeGO on three external datasets. (B) Raw embedding variance distribution (histogram) and PCA explained variance (line) for each dataset and model. (C–E) UMAP comparison of zero-shot, PCA30, and DualStream few-shot embeddings for maize external root (C), rice root (D) and Arabidopsis leaf (E). (F) Quantitative comparison across five metrics (ARI, NMI, inverse Davies-Bouldin, k-NN accuracy, k-NN macro F1) for zero-shot, PCA30 and few-shot methods on each dataset.

Although zero-shot classification accuracy was moderate — consistent with the fact that masked gene modeling (MGM) pretraining optimizes for expression prediction rather than cell-type discrimination — we found that few-shot fine-tuning substantially improved performance. We employed a DualStream architecture that combines raw expression features with CLS embeddings through a lightweight fusion module (∼300K trainable parameters), trained on only 15% of labeled cells from each external dataset. Across all three datasets, few-shot fine-tuning surpassed both zero-shot and PCA30 baselines (Figure 5C–E). On maize root, k-NN F1 improved from 0.292 (zero-shot) to 0.746 (few-shot); on rice root, from 0.398 to 0.894; and on *Arabidopsis* leaf, from 0.553 to 0.926. Clustering metrics including ARI and NMI showed similar trends: on rice, ARI increased from 0.058 (zero-shot) to 0.543 (few-shot), exceeding the PCA30 baseline of 0.211 (Figure 5F). These results indicate that scMaize pretraining provides a strong initialization for representation learning, and that a small amount of labeled data is sufficient to adapt the model for accurate cross-species cell-type classification.

### Encoding of perturbation-condition states beyond cell-type identity

To determine whether scMaize learns cellular states beyond static cell-type identity, we analyzed its embeddings of root cells from three perturbation conditions present in the training atlas: heat stress (PRJCA023597), nitrate treatment (PRJNA759548) and *Fusarium verticillioides* (FV) pathogen treatment (PRJNA865791), totaling 70,959 cells from 7 L2 cell types, with project-matched control cells for comparison (Figure 6). Because treatments were confounded with experimental batches, all analyses used batch-removed embeddings.

**Figure 6.**
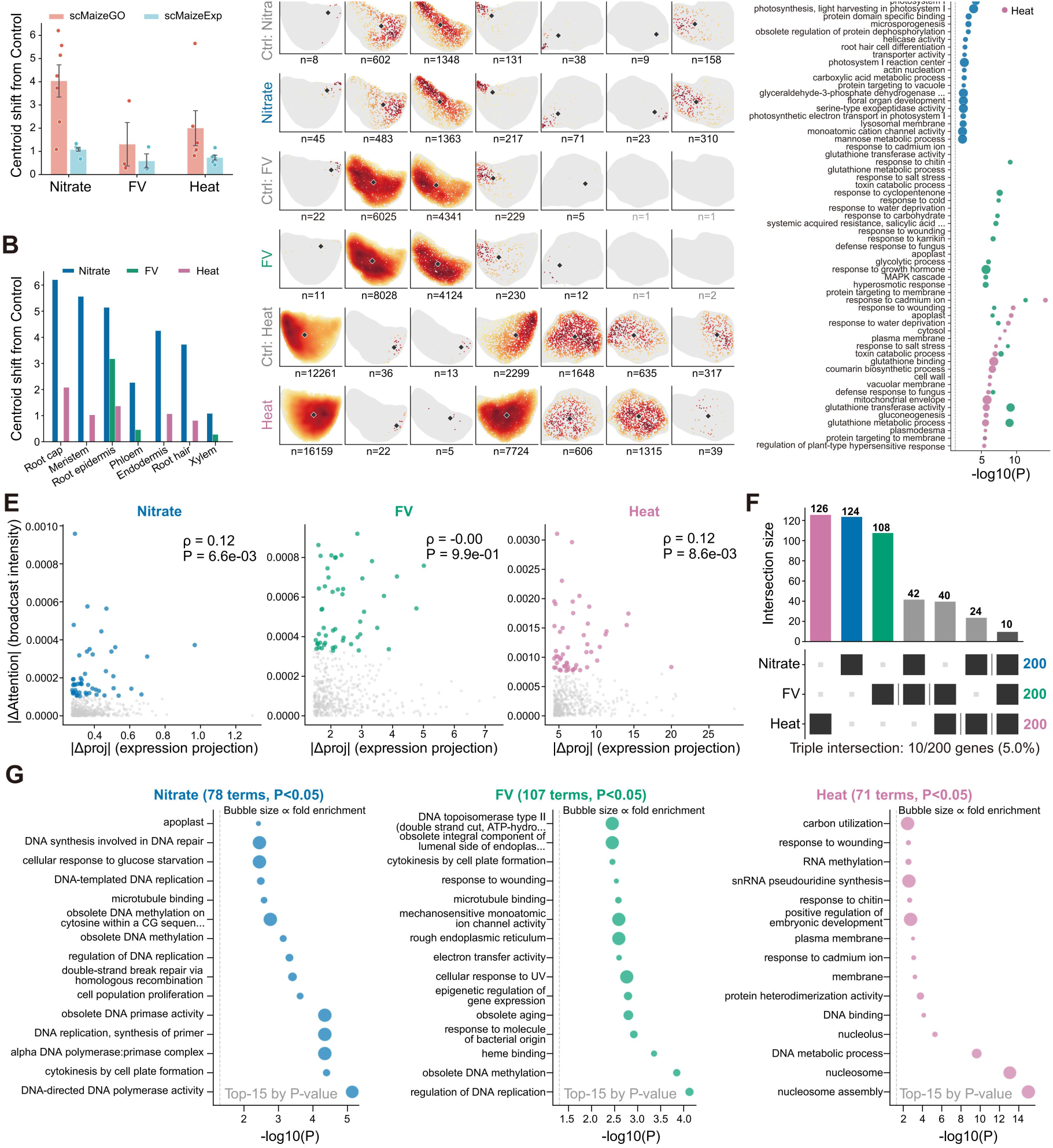
scMaize embeddings encode stress-condition cellular states. (A) Centroid displacement (Euclidean distance in PCA30 space) between treatment and matched control cells for scMaizeGO and scMaizeExp across three stress conditions (mean ± SEM, dots represent cell types, n > 20 per group). (B) Centroid displacement of scMaizeGO per cell type, showing the cell-type-specific hierarchy of stress responses. (C) Density UMAP of 7 root cell types; gray contours denote background density, and diamond markers indicate per-condition centroids. (D) GO enrichment of the top 200 |Δproj| genes per condition (top 20 significant terms, hypergeometric test, P < 0.05), with bubble size indicating fold enrichment. (E) Association between |Δproj| and |Δattn| for 500 genes per condition, with Spearman correlation coefficients. (F) UpSet plot showing intersections among the top 200 genes ranked by |Δattn| across conditions; the P value was obtained from 10,000 permutations of gene-condition labels. (G) GO enrichment of the top 200 genes ranked by |Δattn| per condition (top 20 significant terms, hypergeometric test, P < 0.05), with bubble size indicating fold enrichment.

scMaizeGO produced substantially larger centroid displacements between treatment and matched control cells than scMaizeExp across all conditions (mean 2.76 versus 0.86; Figure 6A), indicating that the GO functional prior amplified perturbation signals ∼3-fold. Centroid displacement varied by cell type and condition (Figure 6B). Under heat stress, root cap showed the highest displacement (2.08), consistent with its role as a protective barrier. Root epidermis, the only cell type with sufficient data across all three conditions, showed the strongest response under nitrate (5.14), followed by FV (3.18) and heat (1.37). Xylem showed the smallest displacements (1.08 and 0.28), consistent with its predominantly dead cell composition, serving as a negative control. Density UMAP visualization across 7 L2 cell types confirmed condition-specific shifts in cellular density distributions, most pronounced in larger cell populations such as root cap and epidermis (Figure 6C).

To identify condition-responsive genes, we quantified treatment–control differences in the model’s expression-projection space (|Δproj|; Supplementary Figure S3). The top-ranked genes showed distinct functional enrichments across conditions (Figure 6D and Supplementary Table S1): nitrate was associated mainly with metabolic and developmental processes, FV with defense and detoxification, and heat with broad stress-response and proteostasis pathways.

Representative genes included ABCG40 and ATTRX under FV, and GSTF3, ABCG40 and UBI10 under heat. Thus, expression projection yielded biologically coherent, condition-associated gene rankings.

Attention changes provided a complementary readout. |Δattn| was only weakly associated with

|Δproj| for nitrate and heat and showed essentially no association under FV (Figure 6E), with no overlap among the top 20 genes from the two rankings within any condition (Supplementary Figure S3A,B). Among the top 200 |Δattn| genes, only 10 were shared across all three conditions, whereas 52–62% were condition specific, and the shared overlap exceeded random expectation (Figure 6F). GO enrichment further revealed distinct functional programs involving cell proliferation under nitrate, DNA replication, methylation and defense-related responses under FV, and chromatin-associated processes under heat (Figure 6G and Supplementary Table S2).

Condition-specific attention hub networks were consistent with these patterns (Supplementary Figure S3C).

### scMaize online platform

To facilitate community access to curated maize single-cell resources, we developed the scMaize web platform (https://www.scmaize.com), which integrates data exploration, interactive visualization, pretrained models and online analysis into a unified interface. Public datasets are processed through metadata curation, quality control, integration and annotation to construct scMaizeAtlas and train the associated models. The current online release comprises 20 projects, 66 samples, 385,675 cells from seven tissues and four platforms, together with 253 downloadable files. The platform web interface is organized into six core pages (Figure 7). The Data page enables browsing and retrieval of project-and sample-level metadata together with associated raw matrices and filtered objects. The Atlas page provides interactive visualization of the integrated scMaizeAtlas through UMAP and t-SNE embeddings, allowing users to explore cell identities, tissue composition and gene expression patterns. The Models page distributes pretrained scMaizeExp and scMaizeGO resources, including model descriptions, performance benchmarks and downloadable weights. The Apps page supports four web-based analysis pipelines—cell-type annotation, embedding extraction, expression imputation and gene similarity analysis—through an asynchronous job management system that accepts user-uploaded datasets and returns downloadable results. In addition, the Download page provides centralized access to atlas data, embeddings and model weights, whereas the Help page offers user documentation, tutorials and contact information, enabling reproducible access to the complete scMaize ecosystem.

**Figure 7.**
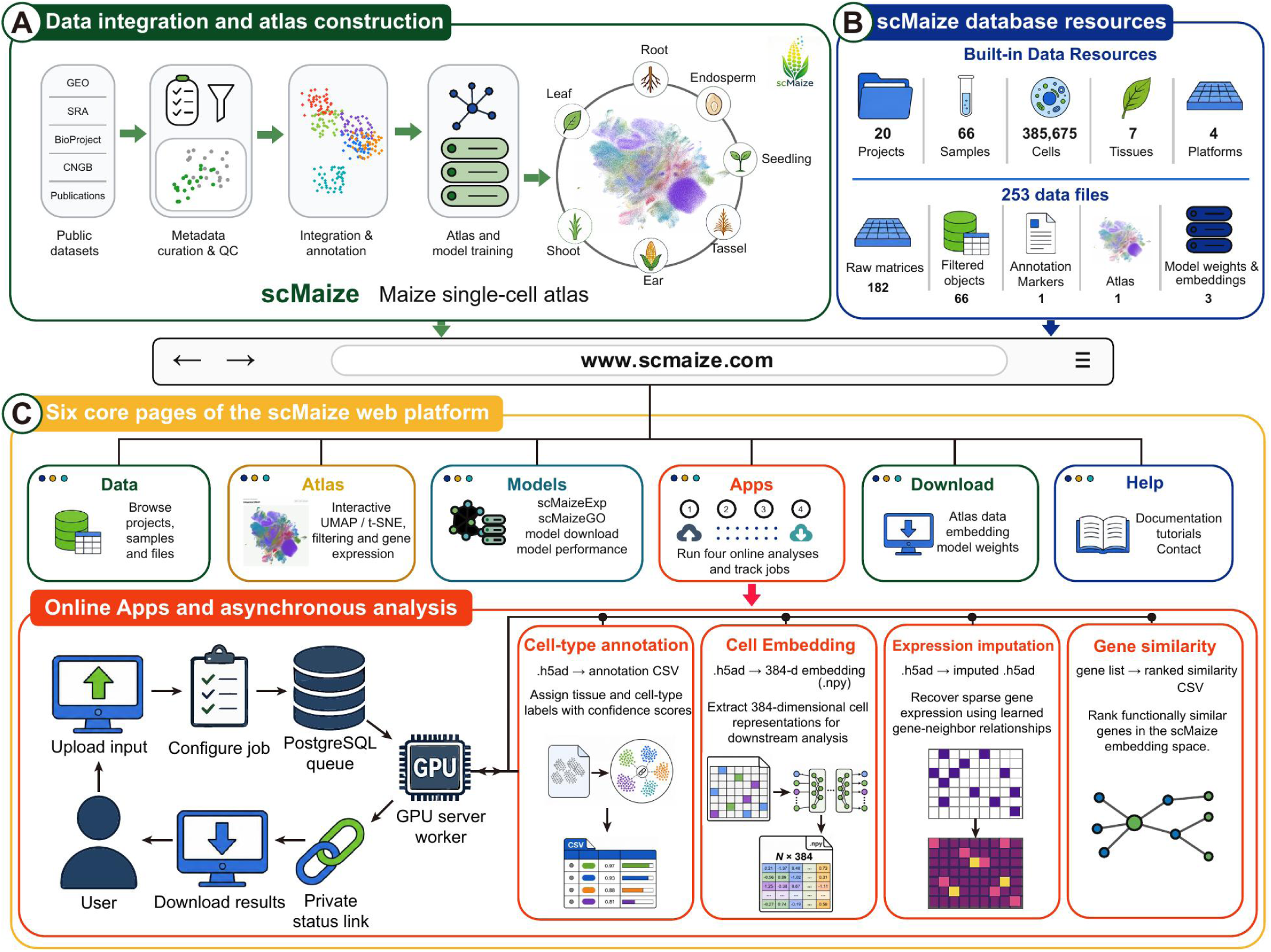
Overview of the scMaize web platform. (A) Construction of the scMaize resource through public data collection, quality control, atlas integration and foundation model training. (B) Summary of the integrated database resources. (C) The six core pages of the scMaize web platform and the asynchronous workflow supporting the four online analysis tools: cell-type annotation, embedding extraction, expression imputation and gene similarity analysis.

## DISCUSSION

scMaize demonstrates that species-specific single-cell foundation models, even at moderate scale, can achieve representations that are accurate within their target organism, transferable across species boundaries, and informative beyond static cell-type identity. The integrated atlas and paired models provide a resource for the maize community, yet the broader contribution lies in the principles they establish for plant single-cell genomics.

A central inference from this study is that species-focused pretraining can be effective at moderate model scale, complementing cross-species strategies such as scPlantLLM and scPlantFormer (Cao et al., 2025; Zhang et al., 2024) that are particularly valuable when species-specific data are scarce. However, our results suggest that species-focused pretraining captures regulatory features that may be diluted in broader cross-taxa representations. A related trade-off has been noted in single-cell modeling more broadly, where domain-specific fine-tuning often unlocks the utility of general-purpose foundation models (Baek et al., 2025). scMaize’s compact architecture (16.7M parameters), trained exclusively on maize data, produced more coherent manifolds and higher cell-type classification accuracy than the larger cross-species scPlantLLM (∼115M parameters). External validation further demonstrates that this species-specific prior transfers across the monocot–dicot divide: zero-shot embeddings generalized to rice (∼50 MYA) and *Arabidopsis* (∼150 MYA), with classification performance primarily reflecting dataset-specific classification difficulty rather than strict phylogenetic distance. While zero-shot classification accuracy was moderate in absolute terms, this is consistent with systematic evaluations of much larger human single-cell foundation models, where zero-shot performance has been shown to underperform even simple baselines (Kedzierska et al., 2025). Notably, few-shot fine-tuning with minimal labeled data (15%) lifted k-NN F1 to 0.746–0.926 across all three datasets, establishing that MGM pretraining captures conserved gene expression programs that transcend species boundaries.

The comparison between scMaizeExp and scMaizeGO demonstrates that functional priors can improve representation learning without materially increasing model parameters. scMaizeExp alone, trained on expression data without GO information, already provides a robust baseline. The addition of GO embeddings did not improve global Pearson correlation for masked expression prediction, but it improved rank-order preservation, strengthened the association between attention and expression similarity, and improved embedding topology. These advantages translated to downstream tasks: scMaizeGO consistently outperformed scMaizeExp across all three external validation datasets, confirming that the GO prior enhances the biological coherence and transferability of learned representations. This pattern, supported by both attention analysis and embedding topology assessments, suggests that GO information contributes less by increasing pointwise prediction accuracy than by regularizing the organization of learned representations — a refinement rather than a replacement of the expression-based foundation. Both models independently highlighted ribosome-associated processes and photosynthesis, consistent with the classical source–sink framework in maize (Guo et al., 2026). Pretraining across cells from both source and sink tissues places these functional modules within a unified cellular representation, which may support future studies of the cellular regulation underlying source–sink coordination.

Structured functional priors, even when incomplete or transferred from model organisms, can serve as effective inductive biases that improve generalization beyond the training distribution. The attention and hub-gene analyses also delineate a boundary of transcriptome-only modeling: both models primarily emphasized highly expressed and co-expressed genes, whereas transcription factors were underrepresented among attention hubs. These observations are consistent with recent work showing that Transformer-based models relying solely on expression data tend to capture co-expression rather than direct regulatory relationships (Kendiukhov, 2026). Moving from co-expression-aware representations toward causal regulatory inference will require multimodal extensions incorporating chromatin accessibility, transcription factor binding, perturbation data or genetic variation (Khan et al., 2026).

Independent of these limitations, the perturbation-condition analysis demonstrates that even a transcriptome-only model can implicitly encode treatment-specific cellular states, despite never being explicitly trained to distinguish conditions. The GO prior amplified perturbation signals approximately 3-fold, and expression projection through the model’s trained weights identified condition-specific gene responses enriched for known stress pathways — defense response to fungus under *Fusarium verticillioides* (FV) treatment, glutathione-mediated detoxification under heat stress, and metabolic reprogramming under nitrate. The expression projection approach is particularly notable because it leverages the model’s internal representations rather than raw expression fold changes. This provides a principled alternative to conventional differential expression analysis, which relies solely on observed expression values and does not exploit learned gene–gene relationships. Attention-based analysis extended this observation by showing that condition-associated changes in gene–gene attention were only weakly associated with expression projection changes and were predominantly condition specific. The absence of a detectable association under FV treatment (ρ ≈ 0) suggests that attention-based and expression-based readouts capture partly distinct aspects of the learned perturbation response. Together, these findings indicate that pretrained model weights contain complementary perturbation information at the expression-projection and attention levels, both of which can be examined without explicit treatment labels.

Several extensions are warranted. Expanding the atlas to additional developmental stages, stress conditions and genetic backgrounds will increase the coverage of maize cellular states. Incorporating epigenomic and spatial modalities should improve cell-type annotation and regulatory interpretation. The DualStream architecture demonstrated that lightweight fusion modules can effectively adapt pretrained representations for supervised tasks; extending this approach to other downstream applications represents a natural next step. Cross-species evaluation beyond the species tested here will be important for understanding the limits of ortholog-based transfer. The online platform lowers the barrier for community adoption.

Taken together, this work establishes the feasibility and value of species-specific single-cell foundation models in plants. The principles demonstrated — that functional priors enhance cross-species transferability, that pretrained representations enable few-shot generalization, and that expression projection enables perturbation-aware gene discovery — extend beyond maize. As single-cell resources for additional crop species continue to expand, the approach of combining unified atlases with species-focused models and structured biological priors offers a scalable template for linking cell-resolved genomics with computational representation learning, with substantial potential to accelerate the application of single-cell technologies to crop improvement.

## MATERIALS AND METHODS

### Data collection and atlas integration

Twenty public maize single-cell RNA-seq projects were identified from NCBI SRA, CNCB GSA and associated publications. Raw sequencing reads were quality-filtered with fastp (v0.23) (Chen et al., 2018). For 10x Genomics libraries, reads were aligned to the maize B73 version 4 reference genome (Jiao et al., 2017), and gene expression was quantified using Cell Ranger (v7.0; 10x Genomics). BD Rhapsody, Singleron and Biomarker datasets were processed according to manufacturer-recommended protocols. Gene expression quantification used the maize B73 version 4 reference annotation. Project-level quality control followed the filtering criteria reported in the original studies and was implemented with Seurat v5 (Hao et al., 2024b). Data processing and integration were performed using OmicVerse (Zeng et al., 2024), with batch correction by scVI (Lopez et al., 2018). Integration quality was evaluated by UMAP visualization before and after correction, with cells colored separately by project and tissue (McInnes et al., 2018).

### Hierarchical cell-type annotation

Marker genes were assembled from scPlantDB (He et al., 2024), PlantscRNAdb (Chen et al., 2021) and manual curation of published maize single-cell studies. Candidate markers were refined through statistical validation, specificity filtering, GO embedding-based deduplication and hierarchical mapping. Differential expression in the integrated atlas was evaluated with the Wilcoxon rank-sum test, and markers with adjusted *P* ≥ 0.05 or log-fold-change < 0.25 were excluded. Markers detected in more than five clusters were removed to reduce broadly expressed genes. GO embedding-based deduplication then merged markers with pairwise cosine similarity > 0.85. Retained markers were capped at 50 per cluster and assigned to broad L2 and fine-grained L3 categories; L3 clusters containing fewer than 100 cells were merged into the corresponding parent L2 category. The final annotation framework comprises 34 L2 and 46 L3 cell types. Annotation confidence was calculated as Σ(score × expression) / marker_count, where score is the Wilcoxon rank-sum statistic; annotation scores were categorized as High (≥0.3), Medium [0.2, 0.3), Low [0.1, 0.2) or Unknown (<0.1) confidence; no cells were assigned to the Unknown category in the final atlas.

### Gene selection and GO embedding

From the 27,754 detected genes, 13,000 highly variable genes (HVGs) were selected using a combined criterion of mean expression and cell-to-cell variance. An additional 2,000 genes were selected iteratively to maximize GO term coverage across Biological Process, Molecular Function and Cellular Component categories. GO annotations were obtained from agriGO (Tian et al., 2017). Four strategies were evaluated for functional embedding construction: IDF-weighted singular value decomposition of GO term assignments (IDF+SVD), ontology-structure embeddings using Onto2Vec (Smaili et al., 2018), BERT-based gene representations using gene descriptions as input text (Devlin et al., 2019), and a hybrid strategy combining IDF+SVD and BERT. Embedding quality was quantified by comparing within-family and between-family cosine similarity across 12 major Pfam protein families (Mistry et al., 2021).

### Model architecture

The scMaize backbone is a 6-layer Transformer encoder (Vaswani et al., 2017) with hidden dimension 384, 4 attention heads and feed-forward dimension 1,536. Dropout was set to 0.1 throughout the model. Each input sequence contains 2,048 gene tokens per cell, randomly sampled from the 15,000-gene vocabulary at each training step and sorted by HVG rank with tight rank encoding. A learnable CLS token is prepended to each sequence and used as the cell-level representation. Four input branches encode gene identity (embedding followed by LayerNorm), expression values (two-layer MLP with GELU activation), batch labels (additive conditioning bias, removable at inference) and, in scMaizeGO only, GO functional embeddings (linear projection). The final CLS output after LayerNorm is used as the 384-dimensional cell embedding. scMaizeExp contains 16.7M parameters and scMaizeGO contains 16.8M parameters.

### Pretraining

Models were pretrained with masked gene modeling (MGM), in which 15% of gene expression values were masked and predicted from the remaining sequence context. A weighted MSE loss applied a 5-fold penalty to non-zero expression values, reducing degenerate convergence toward all-zero predictions in sparse single-cell data. Training used AdamW with learning rate 2 × 10^−4^, weight decay 0.01, 1,000 warmup steps followed by cosine decay, effective batch size 256 (physical batch size 64 with four-step gradient accumulation), and gradient norm clipping at 1.0. Models were trained for 80 epochs on a single NVIDIA A100 40GB GPU, with peak memory use below 15 GB. The 385,675-cell dataset was randomly divided into 80% training, 10% validation and 10% test sets.

### Expression prediction evaluation

Models were evaluated on the MGM task with independent random gene sampling and masking for each cell. Per-gene Pearson *r*, Spearman *ρ* and MSE were computed for genes with at least 10 masked observations. Global metrics were calculated by concatenating predictions across all evaluable genes. Stratified analyses used equal-frequency tertile bins defined by mean expression and expression variance. Per-group medians were accompanied by 95% confidence intervals estimated from 1,000 bootstrap resamples. Kruskal-Wallis tests evaluated differences among strata, whereas paired t-tests and Wilcoxon signed-rank tests were used for cross-model comparisons.

### Attention extraction and analysis

Attention weights were extracted from all 6 Transformer layers for 500 randomly selected training cells using forward hooks. Weights were averaged across layers and heads to obtain one 2,048 × 2,048 attention matrix per cell. The CLS token row and column were removed, and the top-50 target genes per source gene were retained after excluding self-attention. For each cell, we defined *r*_expr_ as the Spearman rank correlation between pairwise gene expression similarity (measured by cosine similarity) and the corresponding attention weights, calculated exclusively on these retained top-50 gene pairs. For cross-model comparisons, paired t-tests and Wilcoxon tests were applied (paired by the same cell across models), with Cohen’s *d* used to estimate effect sizes. Tissue-specific analyses required at least 5 cells per tissue.

### Hub gene identification and functional enrichment

Hub genes were defined as the top-100 genes ranked by cumulative outgoing attention weight across the 500 sampled cells. GO enrichment was tested using Fisher’s exact test against the 2,048-gene HVG background, followed by Benjamini-Hochberg false discovery rate correction. Terms with FDR < 0.05 were considered significant. The 20 most significant terms detected in either model were visualized.

### Cell embedding evaluation

Cell embeddings from 27,365 high-confidence cells were benchmarked against PCA30_HVG, Harmony_HVG (Korsunsky et al., 2019), and scPlantLLM general and maize fine-tuned embeddings (Cao et al., 2025). Classification was performed with K-nearest neighbors (*K* = 30) over five random 80/20 splits. Leiden clustering (Traag et al., 2019) was used for clustering evaluation, with ARI and NMI treated as primary metrics in accordance with recent recommendations to de-emphasize silhouette-based scores in single-cell integration benchmarks (Rautenstrauch and Ohler, 2026). scIB was used to compute iLISI, cLISI, graph connectivity and isolated-label F1 (Luecken et al., 2022). Clustering and integration metrics were computed with Scanpy (Wolf et al., 2018) and scIB, and classification was implemented with scikit-learn (Pedregosa et al., 2011). Qualitative assessment was based on UMAP manifold structure.

### External validation datasets and preprocessing

Three independent single-cell datasets were used for external validation: maize root (PRJCA049374; 19,957 raw and 15,901 post-QC cells), rice root tip (SRP250946; 31,139 raw and 26,138 post-QC cells) and *Arabidopsis* leaf (SRP292306; 5,947 raw and 5,594 post-QC cells). Raw reads were obtained from CNCB GSA for maize and from the NCBI Sequence Read Archive for rice and *Arabidopsis*, and were processed locally following the original study workflows. Rice and *Arabidopsis* cells were annotated after Cell Ranger quantification, Seurat-based quality control and clustering, using canonical marker genes. The maize external dataset was clustered on highly variable genes and annotated by marker expression, with gene identifiers harmonized from RefGen_v5 to RefGen_v4. Cell-type labels from all three datasets were manually matched to scMaizeAtlas L2 categories. Cross-species gene mapping used one-to-one high-confidence orthologs obtained from Ensembl Plants BioMart based on Compara annotations (Tello-Ruiz et al., 2021; Herrero et al., 2016).

### Zero-shot and few-shot evaluation

Zero-shot embeddings were generated using scMaizeExp and scMaizeGO checkpoints with batch embedding removed. Evaluation metrics included k-NN classification accuracy and macro-averaged F1 score (*K* = 30, 5-fold stratified cross-validation), ARI, NMI, Silhouette score, inverse Davies-Bouldin index, and effective embedding dimensions (number of principal components explaining 95% variance). PCA30 on HVG expression was used as the baseline.

For few-shot fine-tuning, a DualStream architecture was employed. The scMaizeGO backbone was frozen, and two trainable projection heads were added: one mapping raw gene expression (2,048 → 128) and another mapping CLS embeddings (384 → 64). The concatenated features (192 dimensions) were processed through a two-layer classifier (192 → 64 → number of classes) with GELU activation and dropout (0.3). Training used 15% of labeled cells from each dataset, with 5% reserved for validation and the remaining 80% held out for testing. Optimization used AdamW with learning rate 1 × 10^−4^, cosine decay with 500 warmup steps, batch size 128, and full FP32 precision. Training ran for up to 50 epochs with early stopping (patience = 20).

### Perturbation-condition analysis

Cells from three root perturbation conditions within scMaizeAtlas were analyzed: heat stress (PRJCA023597, 25,870 treatment and 17,209 control cells), nitrate treatment (PRJNA759548, 2,535 treatment and 2,308 control cells) and *Fusarium verticillioides* (FV) pathogen treatment (PRJNA865791, 12,412 treatment and 10,625 control cells), totaling 70,959 cells from 7 L2 cell types (*n* > 50 cells). Control cells were matched by project and batch to treatment cells. Because treatment identity was confounded with experimental batch, batch embeddings were removed from all CLS representations. For centroid displacement analysis, treatment and matched control cell centroids were computed for each cell-type–condition pair (*n* > 20 per group) in PCA30 space, and Euclidean distance between centroids was reported. Density UMAPs were generated per L2 cell type (n_neighbors = 30, min_dist = 0.3) without subsampling, with control cells on the left and treatment cells on the right. For expression projection analysis, the trained expression projection layer (expr_projection: Linear(1, 384) → GELU → Linear(384, 384)) of scMaizeGO was used to transform per-gene mean expression values into 384-dimensional projected representations. The projection difference |Δ*proj*| = ||proj(treatment_g_) − proj(control_g_)||_2_ was computed for each gene, and the top 200 genes by |Δ*proj*| were subjected to GO enrichment analysis using hypergeometric test against the 15,000-gene vocabulary background, with the top 20 significant terms (*P* < 0.05) reported per condition and all significant terms provided in Supplementary Table S1.

For attention-based perturbation analysis, attention weights were extracted from all 6 Transformer layers of scMaizeGO for 2,000 randomly selected cells in each treatment or matched control group (12,000 cells in total) using forward hooks. Weights were averaged across layers and heads, and self-attention was excluded. For each gene, the attention score was defined as the sum of outgoing weights to all other genes. The attention difference between treatment and matched control was calculated as Δ*attn* = *attn*(treatment) − *attn*(control), and |Δ*attn*| was used for downstream analyses. Spearman rank correlation was used to assess the association between |Δ*proj*| and |Δ*attn*| among the 500 genes included for each condition. Condition specificity was evaluated by intersecting the top 200 genes ranked by |Δ*attn*| across conditions, and the significance of the three-condition overlap was assessed using 10,000 permutations of gene-condition labels. GO enrichment analysis of the top 200 |Δ*attn*| genes was performed against the 15,000-gene vocabulary background using a hypergeometric test (*P* < 0.05; minimum of 5 genes per term). For network visualization, the top 5 genes ranked by |Δ*attn*| in each condition were defined as hubs and connected to their top 3 co-attended neighbors. All significant GO terms are provided in Supplementary Table S2.

### Online platform implementation

The scMaize platform (https://www.scmaize.com) was built as a static website with interactive components. The Atlas module uses a WebGL-based UMAP viewer for cell visualization. The Apps module implements a job queue system for processing user-uploaded single-cell data through the four analysis tools. Model inference on the server uses the same scMaizeGO checkpoint and preprocessing pipeline described above.

## DATA AND CODE AVAILABILITY

Model weights and related data are available at https://www.scmaize.com. All code is available at https://github.com/yjthu/scMaize.

## SUPPLEMENTAL INFORMATION

Supplemental information is available online.

## AUTHOR CONTRIBUTIONS

J.Y. conceived and supervised the project. Q.C. developed the database. Y.Z. collected the data. Q.C., Y.Z., A.Z., and M.S. analyzed the data. Q.C. and Y.Z. drafted the manuscript. J.Y. and X.W. revised the manuscript. All authors reviewed and approved the final version.

## FUNDING

This work was supported by the National Key Research and Development Program of China (2023YFF1000100) and the National Natural Science Foundation of China (32341036).

## DECLARATION OF INTERESTS

The authors declare no competing interests.

## Notes

https://github.com/yjthu/scMaize

https://www.scmaize.com

### Competing Interest Statement

The authors have declared no competing interest.

### Summary of Updates

This revised version expands the perturbation-condition analysis of scMaizeGO. We added a systematic comparison of treatment-associated changes in gene-gene attention and expression projection across nitrate, Fusarium verticillioides, and heat conditions. The new analyses reveal that attention changes are weakly associated with expression-projection changes and are predominantly condition specific. GO enrichment and attention-hub network analyses further show that the two model readouts capture complementary aspects of cellular perturbation responses. Figure 6 was expanded with new panels E-G, and the corresponding Supplementary Figure S3 and Supplementary Table S2 were added or updated. The Abstract, Introduction, Results, Materials and Methods, Discussion, and figure legends were revised accordingly to incorporate these findings and improve the clarity of the manuscript.

