## Supplementary_Figures for "scMaize: A Single-Cell Foundation Model and Integrated Atlas for Maize"

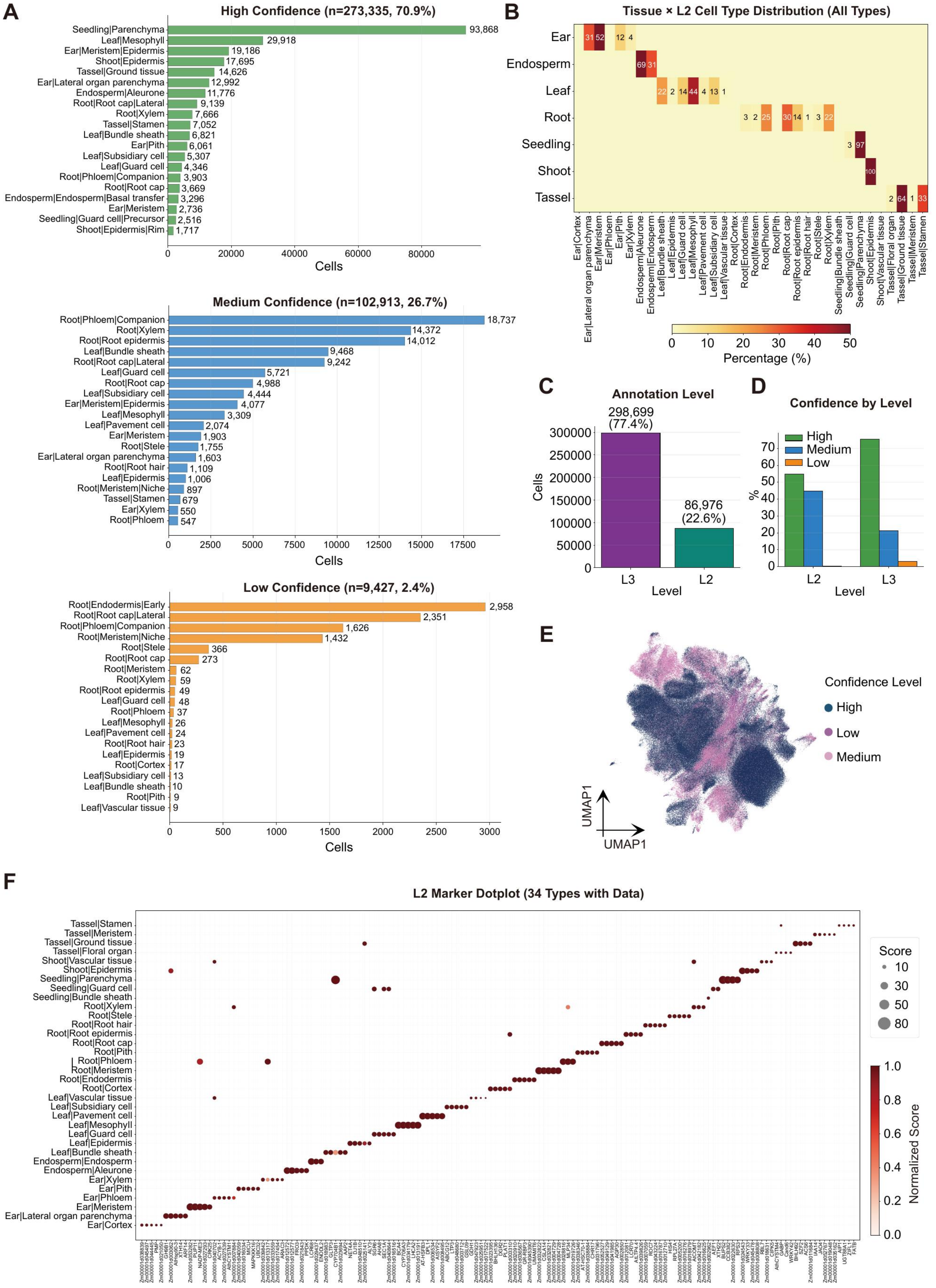


**Supplementary Figure S1.** Annotation statistics of the scMaizeAtlas. (A) Cell-type distributions stratified by annotation confidence level. (B) Tissue composition across annotated cell-type categories. (C) Summary of L2 and L3 annotation coverage. (D) Distribution of High, Medium and Low confidence assignments within the L2 and L3 annotation levels. (E) UMAP colored by annotation confidence. (F) Dot plot of representative L2 marker expression, with dot size indicating the fraction of expressing cells and color intensity indicating normalized mean expression.

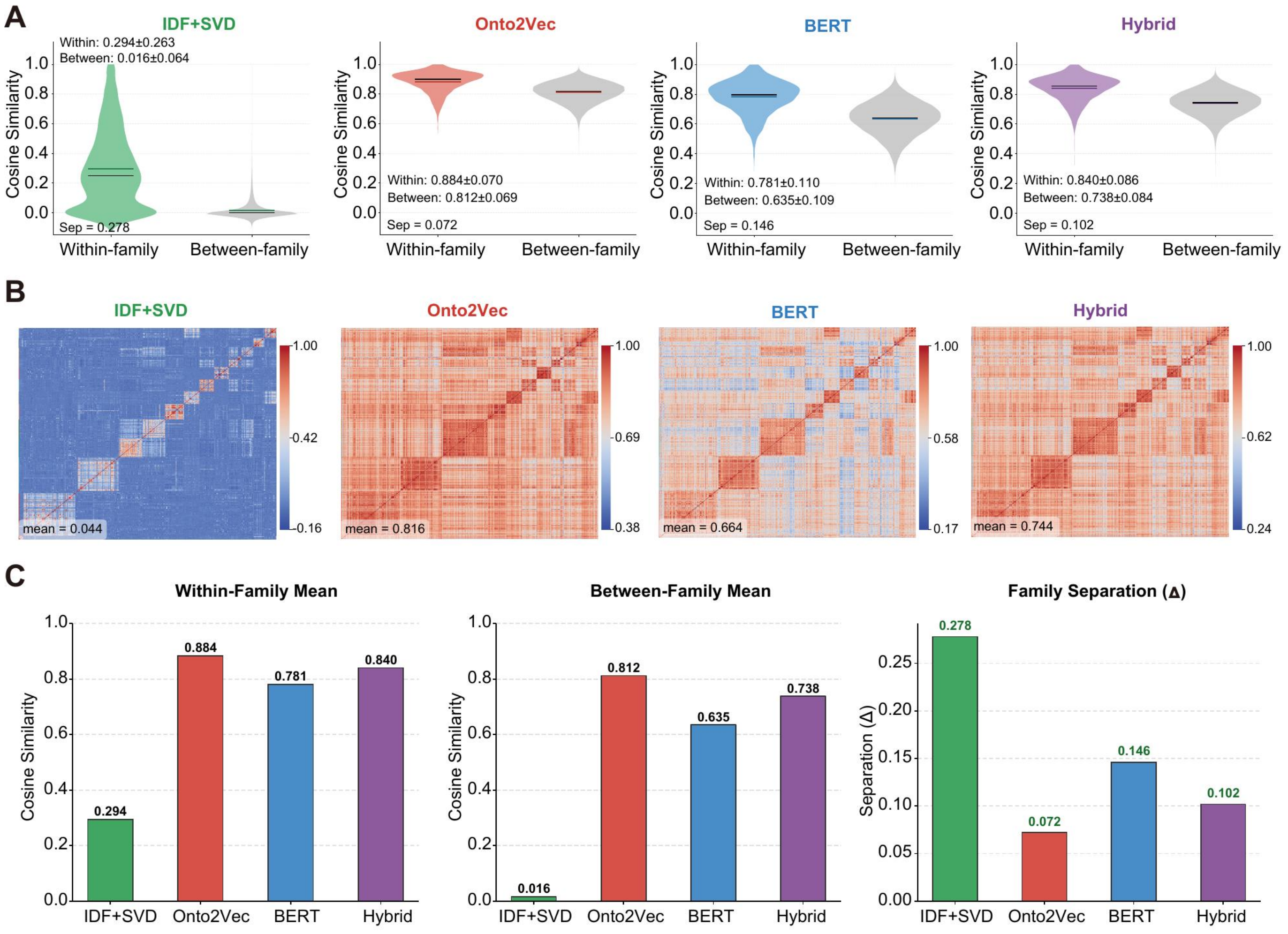


**Supplementary Figure S2.** GO embedding method comparison. (A) Violin plots comparing within-family and between-family similarity across four embedding methods. (B) Gene-gene similarity heatmaps for 12 major Pfam families. (C) Summary of within-family mean, between-family mean and family-separation score for each method.

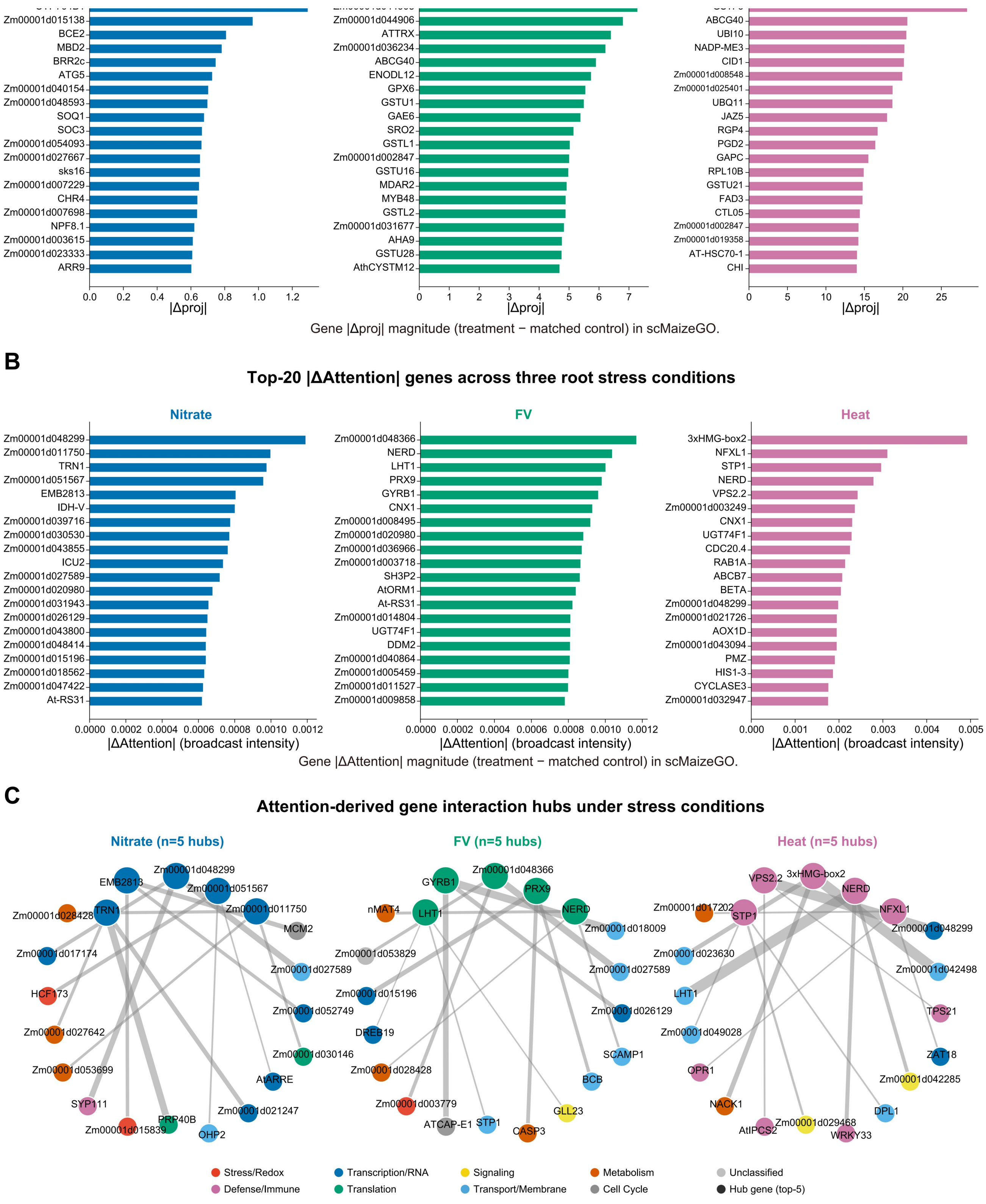


**Supplementary Figure S3.** Expression-projection and attention changes across three root perturbation conditions. (A) Top 20 genes ranked by |Δproj| in each condition. (B) Top 20 genes ranked by |Δattn| in each condition; no genes overlapped between panels A and B within any condition. (C) Attention hub networks showing the top 5 genes ranked by |Δattn| per condition and their top 3 co-attended neighbors. Node color indicates functional category, and unclassified genes are shown in gray.

### SUPPLEMENTARY TABLES

Supplementary Tables S1 and S2 are supplied as separate CSV files.

**Supplementary Table S1.** GO enrichment of expression projection top-200 genes across three root stress conditions. Hypergeometric test results for genes with top 200 $|\Delta proj|$ (Nitrate, 84 terms; FV, 161 terms; Heat, 216 terms; 461 total, $P < 0.05$).

**Supplementary Table S2.** GO enrichment of genes with the largest attention differences across three root perturbation conditions. Hypergeometric test results for the top 200 genes ranked by $|\Delta attn|$ (Nitrate, 78 terms; FV, 107 terms; Heat, 71 terms; 256 total, $P < 0.05$).
